# The dimensionality of plant–plant competition

**DOI:** 10.1101/2021.11.10.467010

**Authors:** Daniel B. Stouffer, Oscar Godoy, Giulio V. Dalla Riva, Margaret M. Mayfield

## Abstract

To avoid extinction, every species must be able to exploit available resources at least as well as the other species in its community. All else being equal, theory predicts that the more distinct the niches of such co-occurring and competing species, the more species that can persist in the long run. However, both theoretical and experimental studies define a priori the nature and number of resources over which species compete. It therefore remains unclear whether or not species in empirically realistic contexts are actually exploiting all or some of the niches available to them. Here we provide a mathematical solution to this long-standing problem. Specifically, we show how to use the interactions between sets of co-occurring plant species to quantify their implied “niche dimensionality”: the effective number of resources over which those species appear to be competing. We then apply this approach to quantify the niche dimensionality of 12 plant assemblages distributed across the globe. Contrary to conventional wisdom, we found that the niche dimensionality in these systems was much lower than the number of competing species. However, two high-resolution experiments also show that changes in the local environment induce a reshuffling of plant’s competitive roles and hence act to increase the assemblages’ effective niche dimensionality. Our results therefore indicate that homogeneous environments are unlikely to be able to maintain high diversity and also shows how environmental variation impacts species’ niches and hence their opportunities for long-term survival.

## Main

Much of our current understanding of species coexistence derives from studying exploitation competition—competition between similar species for a shared pool of finite, limiting resources such as water, nutrients, light, or space^1–6^. In relatively constant environments, the coexistence of many species is thought to depend on two conditions above all others: there should be at least as many resources available as coexisting species^7–9^ and those species should have different niches (e.g., resource-use requirements)^10–13^. When species have distinct niches, intraspecific competition should exceed interspecific competition, preventing communities from becoming overrun by the most dominant competitor or competitors^14,15^. When species are functionally similar, in contrast, their coexistence requires an additional, external source of variability to buffer against otherwise unfavorable years, habitats, or environmental conditions^16,17^.

The theoretical basis linking niche differentiation to coexistence is quite clear^18–21^. The importance of niche differences for maintaining species diversity also has strong empirical support^22–24^. However, we lack a clear picture of the effective number of niche dimensions that actually stabilize population dynamics in diverse natural communities. This is partly due to the fact that researchers traditionally select *a priori* the relevant resources or limiting factors over which species are thought to compete ^8,25^. Unfortunately, such approaches cannot guarantee that all relevant dimensions have been taken into account, or that those selected are even the most relevant ones in practice; even controlled scenarios can therefore overlook previously-unknown, but equally important, mechanisms. Alternatively, researchers have taken a more indirect approach of relating the strength of competition and resource-use variation to differences in functional traits^13^ or evolutionary histories^26^, based on the assumption that these species characteristics are reasonable summaries of the multiple dimensions that compose a species’ niche. Yet indirect, correlative approaches also cannot conclusively identify which resources are limiting^8^ or, more importantly, how many niche dimensions are realized^27^. It therefore remains unclear whether or not the interactions measured for any given ecological assemblage are indicative of highly distinct or highly similar species competing over a large or a small resource pool.

Exhaustive inference of niche dimensionality in diverse empirical communities, as achieved by manipulation of all potential niche axes (e.g., light, water, space, and nutrients) is next to impossible. With this in mind, we introduce here an alternative perspective on this long-standing challenge. Rather than characterize and compare resource use explicitly or select proxies from species’ characteristics, we develop a mathematical and statistical approach that uses information of the strengths and signs of interactions between species to infer their implied niche basis (Methods). In particular, our method provides a means to quantify the “niche dimensionality” of any interacting species assemblage, which is a proxy for the effective number of resources or limiting factors over which those species are competing. From this perspective, niche dimensionality is also a well-defined *mathematical* property of the emergent interactions between species.

In addition to estimating niche dimensionality, our approach decomposes pairwise interactions between species into (i) species’ response traits, (ii) species’ effect traits, and (iii) the relative strength of these responses and effects across the underlying niche dimensions (Fig. 1; Methods). This decomposition helps us identify species’ competitive strategies across shared niche dimensions. Specifically, effect traits modulate the impact *of* each species on all others while response traits modulate the impact of all other species *on* each species^15,28^. When combined, the effect traits of one species and the response traits of another determine whether their directed, pairwise interactions will be strong or weak, competitive or facilitative^26,29^. Given that the net strength of an interaction between two species should be proportional to their niche overlap^7,30^, our framework quantifies species’ niche differences in terms of the extent to which they have an ability to *withstand* neighbor effects (their response) as well as an ability to *generate* neighbor effects (their effect). Each set of response and effect traits thus relates to a comparable set of “effective resources” such that a species with a large effect trait can be thought of as one that depletes the corresponding resource to the detriment of others, and a species with a large response trait can be thought of as one that is particularly sensitive to scarcity of that same resource. As such, response and effect traits allow us to capture the multi-dimensional “strategies” employed by species to outcompete each other or to avoid being outcompeted^27,28^ based directly on the observed outcomes of species-species interactions (as opposed to indirectly; e.g., via their phenotypic traits^31^).

**Figure 1:**
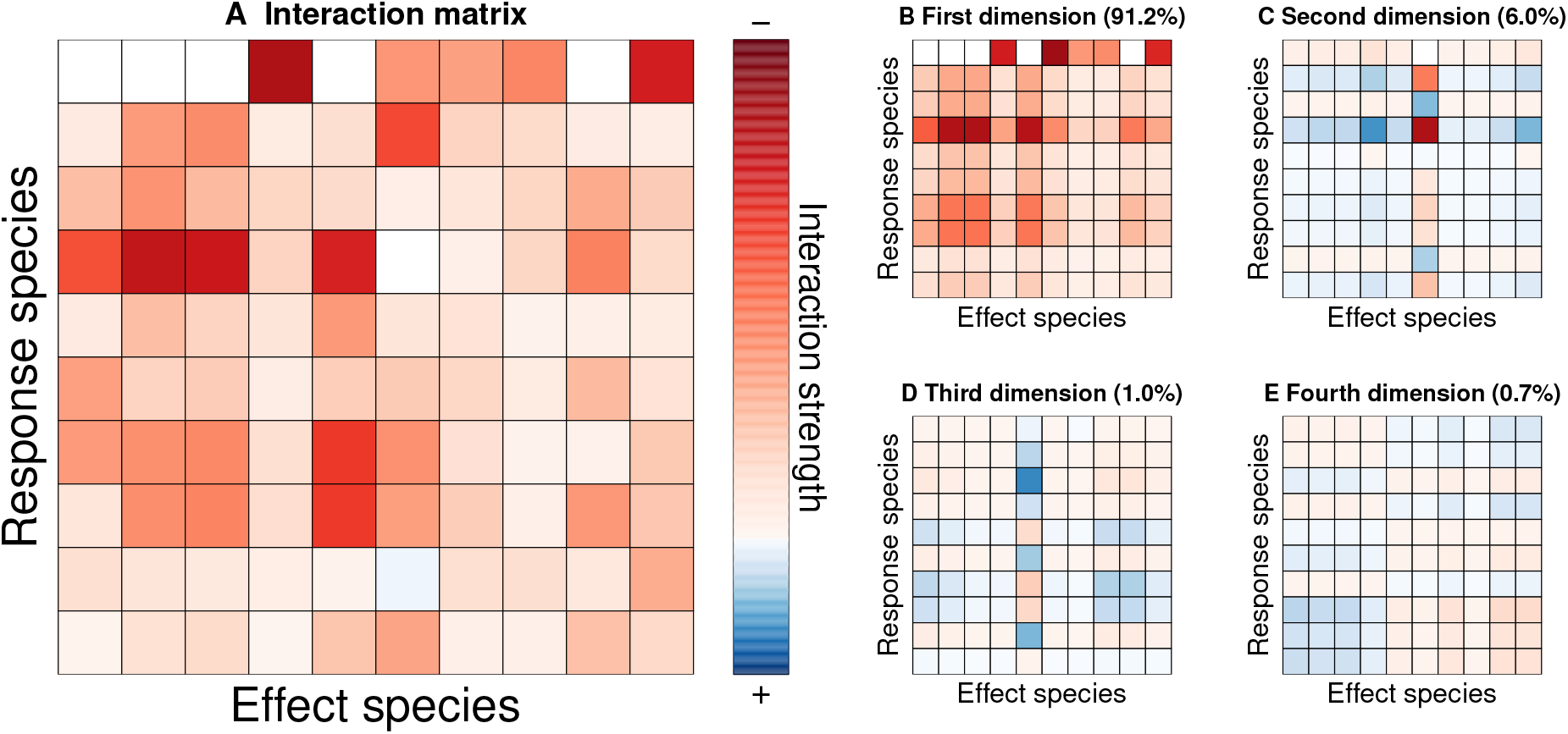
Response-effect decomposition of a pairwise interaction matrix. **A**, The inferred 10 × 10 interaction matrix for Dataset 1-Wet, where rows indicate the species “responding” to the interaction and the columns represent the species “effecting” the interaction. Interactions are colored based on the strength and sign of the net pairwise impact: net competitive values are indicated from white to red while net facilitative values are indicated from white to blue. **B, C, D**, and **E**, The full pairwise interaction matrix can be parsimoniously decomposed into matrices that capture implicit niche differentiation underpinning interactions across the assemblage. For this dataset, the first, second, third, and fourth niche dimensions explain 91.2%, 6.0%, 1.0%, and 0.7% of the variation observed in the data, respectively. Note that interactions in the first niche dimension are overwhelmingly competitive whereas species-specific variation within the second through fourth dimensions both strengthen and weaken the net strength of pairwise interactions.

To investigate whether there are common patterns of niche dimensionality across ecological communities, we applied our approach to 12 empirical assemblages drawn from plant communities across the globe (Methods). These assemblages cover a broad array of habitat types and plant life-history strategies, from deserts to forests and annual plants to trees (Supplementary Information). Each of these includes data regarding the strength and sign of species interactions between three to ten different plant species, and allows us to determine whether each species performs better or worse in the presence of all others. Moreover, the data come from field or common garden settings—in which we might expect extrinsic factors could give rise to greater realized variation—and greenhouses—in which we might expect variation to be driven almost entirely by intrinsic species properties.

## Results

Given these 12 assemblages, we first determined their niche dimensionality 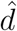: the minimum number of niche dimensions required to explain the observed variation within the data (Methods). As noted above, niche dimensionality 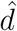 gives an indication of the effective number of resources over which the species in these different plant assemblages are likely competing, and hence the number of species predicted to coexist in the absence of exogenous variation. For every dataset and regardless of the experimental context, we observed that three or fewer niche dimensions 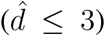 were sufficient to accurately capture the pairwise interactions between species (Fig. 2 & Figs. S1–S3). In fact, the first niche dimension alone explained on average 86.7% of the variation in the plant–plant interactions, and this ranged from 59.6% to 99.3% across the 12 datasets. To verify that these small values of 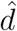 are an ecological feature of these species assemblages and not an artifact of our methodology, we also estimated the niche dimensionality for simulated data with randomly-assigned interaction coefficients. As expected, randomization tends to create “unstructured” data with niche dimensionality closer to the number of species and hence greater than observed in the natural and experimental assemblages (Supplementary Information).

**Figure 2:**
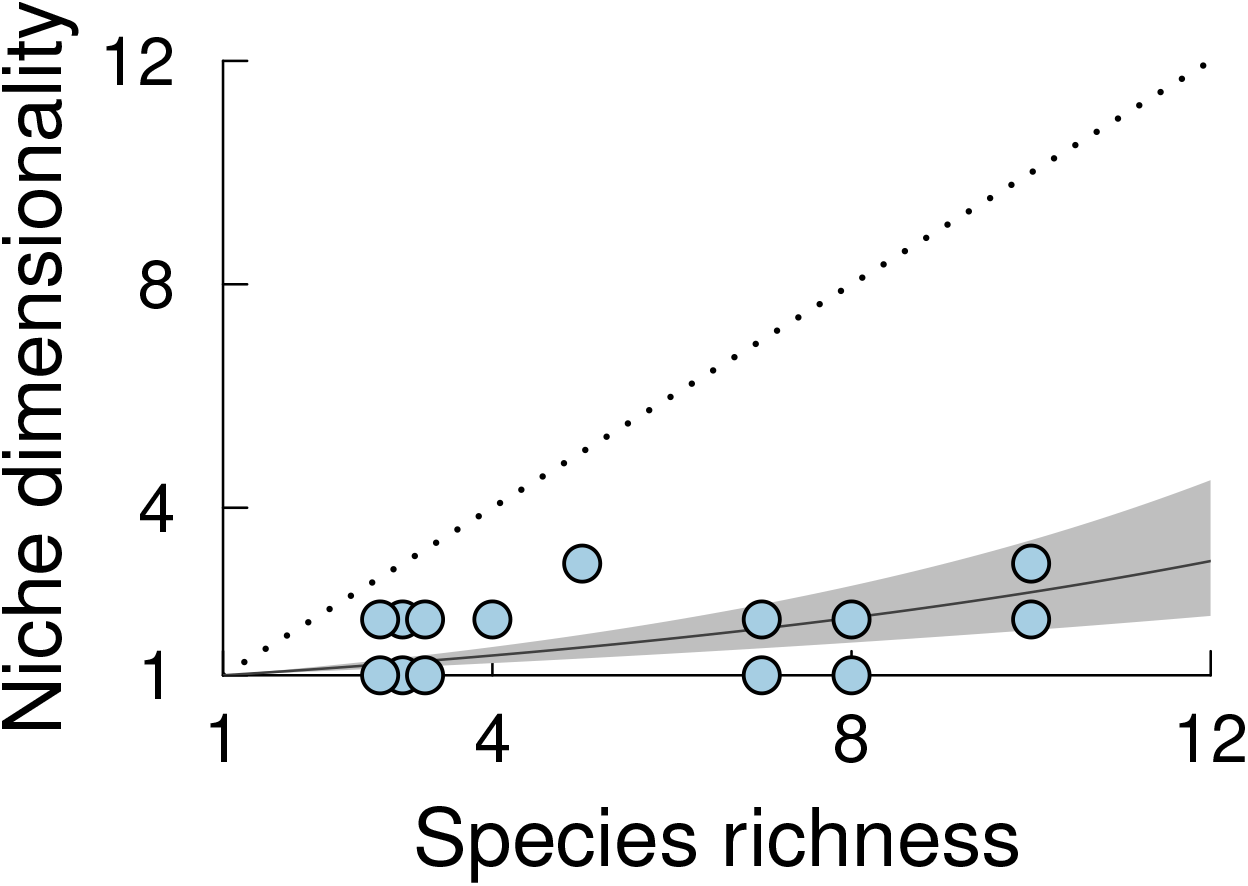
Inferred niche dimensionality 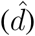 relative to the actual species richness of each empirical assemblage (*S*). The dotted line represents the upper bound of niche dimensionality equal to the number of species whereas the solid line represents the predicted increase of niche dimensions (± standard error) based on the trend observed across the empirical datasets. Note that datasets with identical values of 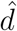 and *S* have been jittered slightly to make them visible.

After estimating niche dimensionality of each dataset, we next examined species’ inferred response and effect traits to get a clearer picture of the forces underpinning species interactions. As representative examples, the best-fit parameters for Dataset 1-Wet and Dataset 1-Dry indicate that the 10 constituent species play distinct ecological roles across their leading niche dimensions (Fig. 3). In both cases, species are sorted into “classic” competitive hierarchies within which most species with strong effects tended to have weak responses and those with weak effects tended to have strong responses. However, realized niche differentiation in Dataset 1-Wet was driven by greater variation in species’ effects than their responses whereas species had heterogeneous responses and heterogeneous effects in Dataset 1-Dry. We observed similar patterns across each of the empirical datasets (Fig. S4), indicating that the leading dimensions of niche differentiation are those that create variation in species’ responses to and effects on other species in their community.

**Figure 3:**
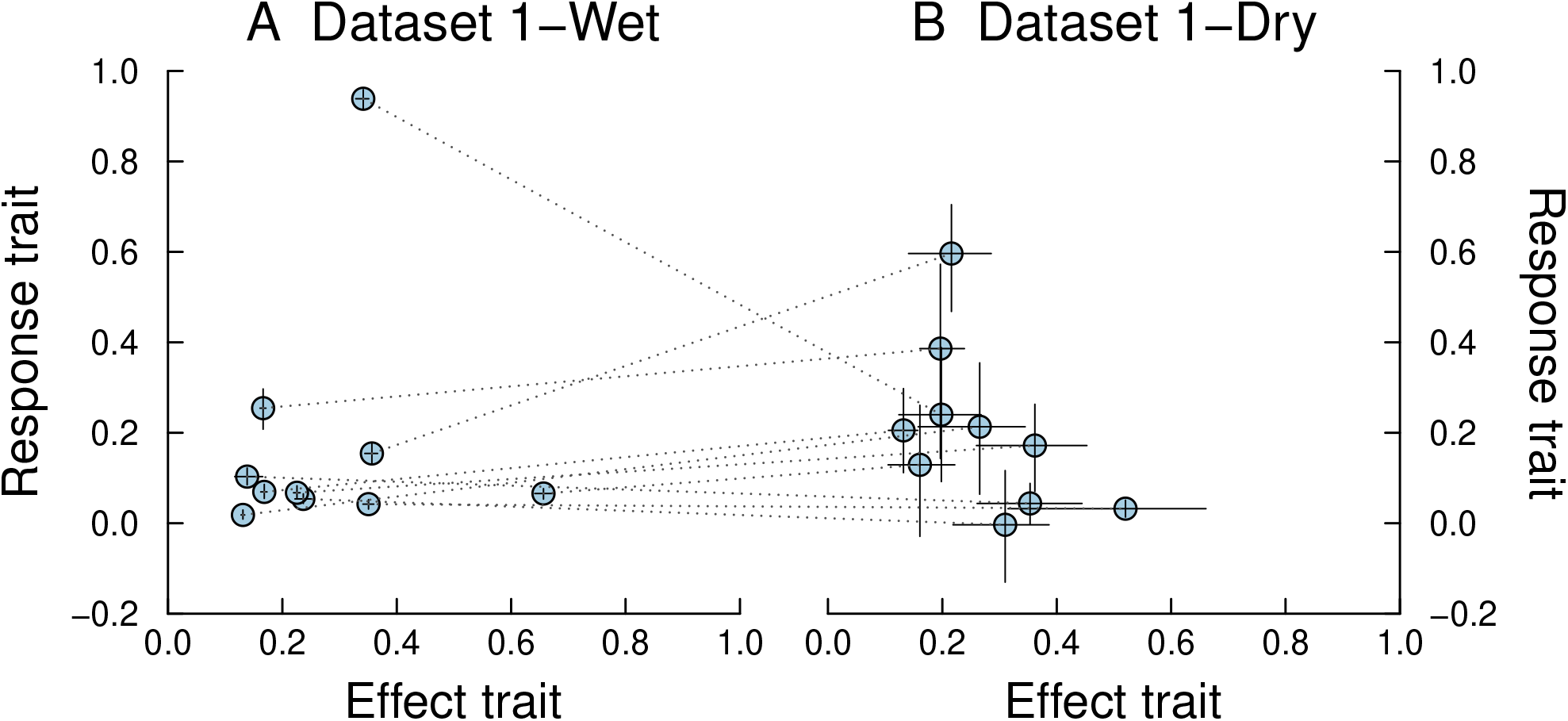
Species’ response and effect traits in the first niche dimension for both Dataset 1-Wet and Dataset 1-Dry. **A**, In Dataset 1-Wet, species are organized along a competitive hierarchy driven largely by variation in their effects as neighbors: dominant competitors have large positive effect traits and small positive response traits whereas weaker competitors have small positive effect traits and large positive response traits. One notable species clusters outside this hierarchy by exhibiting a moderate effect trait and large positive response trait, indicating that it is particularly susceptible to competitive effects. **B**, In Dataset 2-Dry, species are again organized in a competitive hierarchy but exhibiting clearer variation in both response and effect traits. The dotted lines connect species’ response and effect traits in Wet environmental conditions to those same species’ response and effect traits in Dry environmental conditions. Variation between where species fall in the two panels is thus indicative of reorganization of the underlying competitive hierarchy. In both panels, the error bars at each point indicate the 25th to 75th percentile confidence interval about their inferred values and have been plotted on top of points to facilitative visibility when they are small.

We consistently found that estimates of niche dimensionality 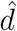 were lower than the total number of species *S* in each empirical assemblage (Fig. 2), and this did not depend on whether or not data came from experiments in the field, garden, or greenhouse (Table S1). Though niche dimensionality increased with increasing species richness (Fig. 2), the rate of increase was much lower than one dimension per species as expected from theory^7–11^. When niche dimensionality is less than the number of species, the interactions of two or more species directly mirror each other; should any of these functionally-similar species have even the slightest competitive advantage, it will tend to dominate in the long run^1,7^. Our results therefore are indicative of communities that can sustain limited diversity, despite bountiful evidence to the contrary in nature. Since some other mechanism must therefore be at play, we next determined whether changing environmental conditions increases niche dimensionality and hence the prospect of coexistence. To do so, we explored two of our datasets in greater detail as they each comprise the same underlying assemblages interacting in two distinct environments. One pair of datasets (Dataset 1, Wet & Dry) consists of ten species in control and simulated-drought conditions^32^, designed to mimic the impact of extreme climatic events of the sort expected with ongoing global change^33^; the other pair of datasets (Dataset 2, Sun & Shade) consists of eight species in control and artificially-shaded conditions, since light availability is known to structure local diversity in that system^34^.

Viewed independently, Dataset 1 was captured by two and three niche dimensions in the Wet and Dry environments while Dataset 2 was captured by one and two niche dimensions in the Sun and Shade environments (Table S1). Nevertheless, the question remains whether the species in each Dataset sorted themselves along the *same* leading axes of niche variation in the contrasting environments. To check this, we measured the correlation between species’ response and effect traits inferred under the two environmental conditions. These correlations were weak and non-significant (Procrustes *ρ* = 0.52, *p* = 0.37 for Dataset 1; Procrustes *ρ* = 0.40, *p* = 0.06 for Dataset 2), indicating that niche dimensions differ depending on environmental conditions. Environmental variation therefore leads to significant shifts in species’ absolute and relative niches (Fig. 3), in contrast to niche theories based on environmental tolerance^35^ or functional traits^36^ which generally assume that niches are immutable species’ attributes.

## Discussion

Much research is dedicated to exploring how the abiotic environment impacts species’ performance in an *interaction-free* context^37,38^. Indeed, most predictions about how communities will change in the face of external disturbance, such as climate variability, depend on this relationship^33,35^. Comparatively less is known about how environmental conditions impact the pairwise interactions in multi-species assemblages and therefore about species’ ability to persist under changing environmental conditions^32,39,40^. One possible reason is that it is difficult to relate changes across multivariate interactions between species to changes in univariate single-species outcomes^21,41^. Here, we show how to decompose pairwise interactions into species-specific response and effect traits, thereby simplifying such inquiries. Moreover, the effect of the environment on species’ response and effect traits generates multiple ecologically-relevant ways for pairwise interactions to vary from one environment to another: all else being equal, any species whose response traits increase in value from one environment to another will face harsher competition; in contrast, any species whose effect traits increase in value from one environment to another should become increasingly dominant. In addition to demonstrating that low-dimensional competition is pervasive in plant assemblages, our approach provides a new lens through which to interrogate coexistence in varying environments.

Despite the differences between the datasets we have studied—such as their biogeographical provenance, evolutionary histories, or growth form—we provide unambiguous evidence that plant–plant interactions are organized over a small number of effective niche dimensions, contrary to common theoretical expectations. While this fact would also appear to imply limited prospects for coexistence, low niche dimensionality in terms of species’ interactive responses and effects is in strong agreement with other assessments of global plant diversity being captured by a small number of life histories^27^ and limited phenotypic trait combinations^36,42^. It also agrees with numerous studies that found that models parameterized with empirical data rarely predict coexistence^13,41,43^. And yet the environment leaves such a strong imprint on interactions that its variability generates one to many additional niche dimensions and serves as a de facto “landscape” across which species can evolve novel strategies to avoid being outcompeted^23,27,44^. While this can make it harder to identify consistent structure in empirical interaction matrices^45,46^, it also highlights a critical link between changing environmental conditions and species’ realized niches^44,47^.

## Methods

### Empirical data

We analyze 12 empirical assemblages to test our core hypothesis regarding variation in the structure of pairwise interactions between co-occurring plants (Table S1). For Datasets 1-Wet, 1-Dry, 2-Sun, and 2-Shade, we directly analyzed raw empirical data from experimental studies of competition between annual plants in order to estimate interaction strengths. For Datasets 3–12, we analyzed previously-estimated pairwise interaction matrices that were available in the literature.

### Estimating interaction strengths with fixed niche dimensionality

When constrained to a fixed dimensionality *d*, our mathematical approach separates every pairwise interaction *α*_*ij*_ into *d* separate components which we refer to as “niche dimensions”. For the simplest case of *d* = 1, the per capita strength of the effect of species *j* on species *i* is given by *α*_*ij*|*d*=1_ = *σ*_1_ × *r*_*i*,1_ × *e*_*j*,1_; that is, by the product of the average strength of interactions in the first dimension (*σ*_1_), the “response trait” of species *i* in the first dimension (*r*_*i*,1_), and the “effect trait” of species *j* in the first dimension (*e*_*j*,1_). As described in the main text, we refer to the *r*_*i,k*_ parameters as responses because they influence how the performance of the same focal species *i* “responds” to the presence of different neighbor plants; similarly, we refer to the *e*_*j,k*_ parameters as effects because they influence how the same neighbor plant “effects” the performances of different focal plants. The separation of responses from effects allows us to generate asymmetric interaction matrices—where the effect of species *i* on *j* differs from the effect of species *j* on *i*—which is more empirically realistic^45^ than the symmetric counterpart commonly studied in theoretical contexts^48,49^. For any value of *d* > 1, the net strength of a pairwise interaction between *i* and *j* is given by 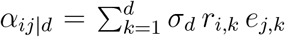, where the sum is across the different niche dimensions and the average strength of these dimensions is ordered such that *σ*_1_ > *σ*_2_ > … > *σ*_*d*_. In this way, the net strength of each interaction is additively partitioned across niche dimensions.

### Density-dependent plant performance

Each of Datasets 1-Wet, 1-Dry, 2-Sun, and 2-Shade consists of estimates of individual plant performance and the abundance or density of co-occurring plants within interaction neighborhoods. To infer interaction strengths, we therefore first had to define a mathematical model for how performance varies as a function of neighbor composition and abundance. In line with current best practice^50^, we estimated the per capita effects of neighboring species on the performance of focal individuals of each of the datasets using a model of the form

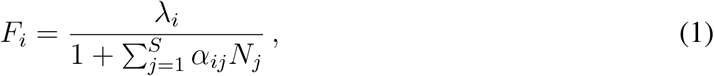

where *F*_*i*_ is the observed estimate of the performance of a focal individual from species *i, λ*_*i*_ captures intrinsic performance of these individuals in the absence of competition, *α*_*ij*_ is the per capita impact of species *j* on species *i, N*_*j*_ is the abundance of species *j* in the focal individual’s interaction neighborhood, and the sum is across all species in that neighborhood (which could potentially include conspecifics of species *i*). For an assemblage of *S* species, our goal then was to estimate the *S* × *S* pairwise interaction matrix *A*. Given data of focal-plant performance and those plants’ interaction neighborhoods, inference of both intrinsic performance and the strength of pairwise interactions can usually be achieved with standard regression approaches^51^. However, these are no longer feasible when constraining interactions to occur at a fixed niche dimensionality. We therefore inferred the parameters of our model by exploiting the similarity of this mathematical model of competition^26^ to singular value decomposition (*SVD*)^52^. To further facilitate parameter estimation, we constrained the inferred response and effect traits so that they always create orthogonal matrices. As a result, we inferred *d*×(2*S* − *d*) total parameters for the interactions at any given niche dimensionality *d*.

### Inferring best-fit reduced-dimensionality model parameters

When we only had an empirical interaction matrix *A* for a given assemblage, we found the maximum-likelihood reduced-dimensionality representation directly using singular value decomposition (*SVD*)^52^. Specifically, one can factorize *A* into three separate matrices such that *A* = *R*Σ*E*^*t*^. Here, *R* and *E* are *S* × *S* orthogonal matrices (i.e., matrices whose columns and rows are all orthogonal unit vectors), and Σ is a matrix with the singular values of *A* along the diagonal and zeroes elsewhere. We can equivalently factorize the matrix as 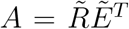 by defining 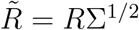 and 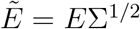 and where Σ^1/2^ is the Cholesky decomposition of Σ. *SVD* works in such a way that 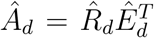 is the best least-squares approximation of *A* at any resource dimensionality *d* ≤ *S* as long as 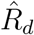 is the *S* × *d* sub-matrix given by the first *d* columns of 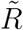 and *Ê*_*d*_ is the *S* × *d* sub-matrix given by the first *d* columns of 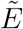. As a consequence, the maximum likelihood values of the response and effect parameters in the first *d* resource dimensions correspond to the first *d* columns of *R* and *E*, respectively.

When we had raw empirical data and hence the values composing the interaction *A* were unknown, we inferred them from the raw empirical data. For any fixed resource dimensionality *d*, this required estimating the *d* values along the diagonal of Σ_*d*_ and the two *S* × *d* matrices *R*_*d*_ and *E*_*d*_. To maintain coherence with *SVD*, the columns of *R*_*d*_ were maintained to be orthogonal unit vectors. This implies that the first column has *S* − 1 free parameters, the second column has *S* − 2 free parameters, and so on, up to a maximum of *S* (*S* − 1) /2 free parameters if *d* = *S*. The same is true for the columns of *E*_*d*_. In total, we must therefore perform inference on *d* × (2*S* − *d*) parameters. In order to ensure that the parameters of *R*_*d*_ and *E*_*d*_ maintain the orthogonality constraint and uniqueness during optimization, we followed the Cayley transformation method of Shepard et al.^53^ and Jauch et al.^54^.

We used the *mle2* function from the *bbmle* package^55^ in the statistical programming language R ^56^ to identify the optimal values of Σ_*d*_, *R*_*d*_, and *E*_*d*_. Conveniently, Eq. (1) can be fit as a Poisson regression with inverse link function^57^. To facilitate comparison to the approach with which the matrix *A* is not factorized into its response–effect equivalent, we therefore used *mle2* to minimize the exact same deviance function as used for Poisson regression rather than to maximize the data’s log-likelihood.

### Statistical analysis

For Datasets 1-Wet, 1-Dry, 2-Sun, and 2-Shade, we inferred the best-fit parameters for interactions constrained to occur when dimensionality *d* = *S*. We then used the inferred values of 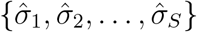 and identified the niche dimensionality 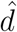 that was sufficient to capture 95% of the variation in the inferred interaction matrix. For Datasets 3–12—for which we only had an estimated interaction matrix *A*—we determined the niche dimensionality 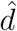 based directly on the singular values of *A*^52^. For all dataset types, values 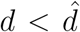 fail to capture biologically meaningful variation in the observed plant–plant interactions; values of 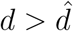 require the use of an overly complex statistical model with an excess of resource dimensions. Though we expect that values 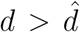 will give a more refined description of species’ fundamental niches, 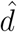 is a better reflection of the niche differences that are *actually realized* as a result of species–species interactions.

We then studied the properties of the 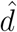-dimensional estimates of species’ response and effect traits within each dataset. For the paired Datasets 1 & 2, we tested whether the positions of species in each of these 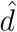-dimensional spaces were correlated using a Procrustes analysis. A significant Procrustes statistic indicates that the effective response and effect “hierarchies” between species in both environments are positively related (e.g., species always tend to be the strongest or weakest effectors or responders) whereas a non-significant result implies that species likely play different roles when the environmental conditions change.

## Code availability

The code used for analyses in this study can be found at https://github.com/stoufferlab/dimensionality-of-competition.

## Data availability

All data used in this study can be obtained from https://doi.org/10.5061/dryad.8v13t2q, https://doi.org/10.5061/dryad.5d1s9, https://doi.org/10.5061/dryad.1sm06sp, or the original references as listed in Table S1.

## Acknowledgements

We thank Bernat Bramon Mora, Hao Ran Lai, and Anne McLeod for providing comments and suggestions on the manuscript. We acknowledge the support of a Rutherford Discovery Fellowship (RDF-13-UOC-003 to DBS) and a Marsden Fund Grant (16-UOC-008 to DBS and MMM), both administered by the Royal Society New Zealand Te Apārangi from NZ Government funding, an ARC Discovery Grant (DP140100574 to MMM and DBS), and a Ramón y Cajal Fellowship provided by the Spanish Ministry of Economy and Competitiveness (MINECO) and by the European Social Fund (RYC-2017-23666 to OG).

