## Supplementary Information for "The dimensionality of plant–plant competition"

**Table S1:** Dataset details and inferred niche dimensionality. For each plant assemblage, we show its Country of origin, the Habitat the species are drawn from, the Experimental setting, the corresponding Environmental treatment (if applicable), whether we have empirical data with which to fit a model for density-dependent performance (Raw) or a matrix of interaction coefficients/relative performance (Matrix), the Species richness $S$, the inferred Niche dimensionality $\hat{d}$, and the original Reference within which to find all experimental protocols.

| Dataset | Country | Habitat | Experimental setting | Environmental treatment | Data format | Species richness ($S$) | Niche dimensionality ($\hat{d}$) | Reference |
|---|---|---|---|---|---|---|---|---|
| 1 | ESP | Grassland | Field | Wet | Raw | 10 | 2 | Matías *et al.*[1] |
| 1 | ESP | Grassland | Field | Dry | Raw | 10 | 3 | Matías *et al.*[1] |
| 2 | AUS | Open woodland | Field | Sun | Raw | 8 | 1 | Stouffer *et al.*[2] |
| 2 | AUS | Open woodland | Field | Shade | Raw | 8 | 2 | Stouffer *et al.*[2] |
| 3 | USA | Old field | Greenhouse | - | Matrix | 7 | 1 | Goldberg & Landa[3] |
| 4 | USA | Shrubland | Garden | - | Matrix | 7 | 2 | Kinlock[4] |
| 5 | PAN | Tropical forest | Field | - | Matrix | 5 | 3 | Svenning *et al.*[5] |
| 6 | CHN | Desert | Greenhouse | - | Matrix | 4 | 2 | Gao *et al.*[6] |
| 7 | BRA | Estuarine | Field | - | Matrix | 3 | 2 | Costa *et al.*[7] |
| 8 | ESP | Old field | Garden | - | Matrix | 3 | 2 | Domènech & Vilà[8] |
| 9 | USA | Grassland | Field | - | Matrix | 3 | 2 | Farrer & Goldberg[9] |
| 10 | USA | Old field | Greenhouse | - | Matrix | 3 | 1 | Gurevitch *et al.*[10] |
| 11 | USA | Grassland | Greenhouse | - | Matrix | 3 | 1 | Hedberg *et al.*[11] |
| 12 | GER | Grassland | Garden | - | Matrix | 3 | 1 | Weigelt *et al.*[12] |

#### Niche dimensionality of empirical datasets

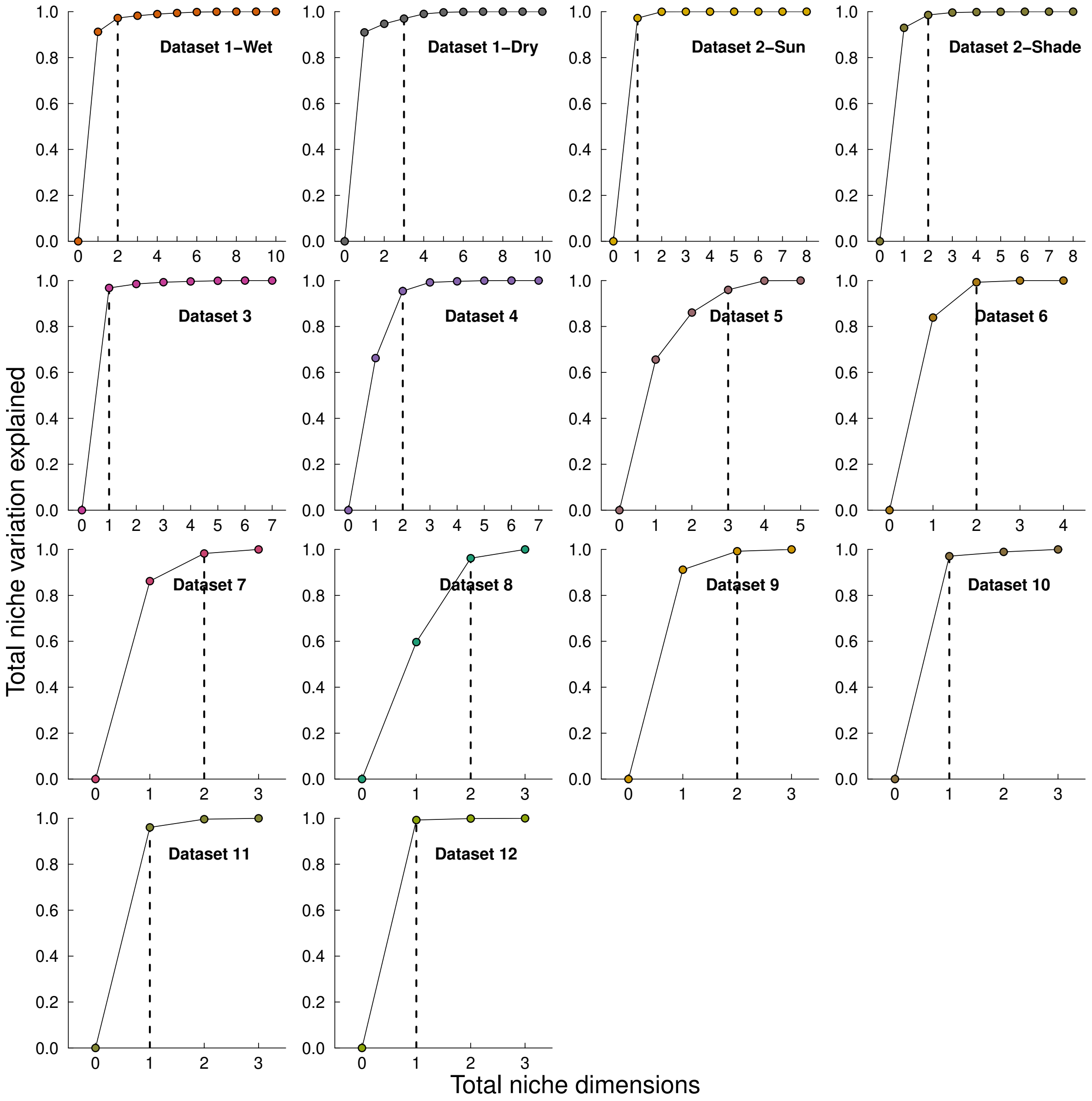


**Figure S1:** For each matrix-only dataset, we show the total variance explained when interactions are inferred at a fixed number of niche dimensions. The dashed line indicates the inferred niche dimensionality $\hat{d}$ required to explain at least 95% of the variation in the observed interactions.

#### Niche dimensionality of empirical datasets (continued)

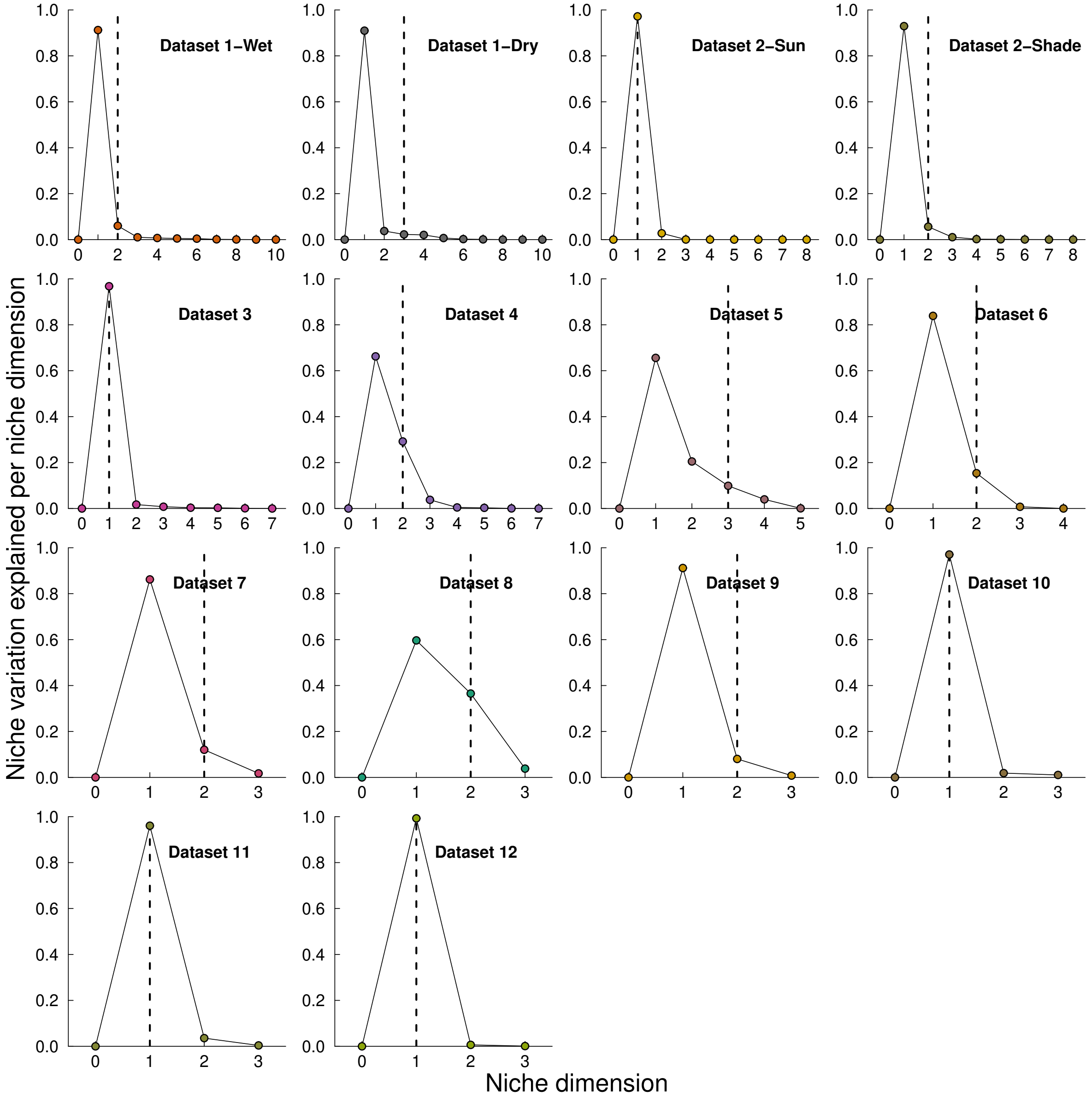


**Figure S2:** For each matrix-only dataset, we show the incremental variance explained by each niche dimension when interactions are inferred at niche dimensionality $d = S$. The dashed line indicates the inferred niche dimensionality $\hat{d}$ required to explain at least 95% of the variation in the observed interactions.

#### Observed versus fit interactions for all datasets

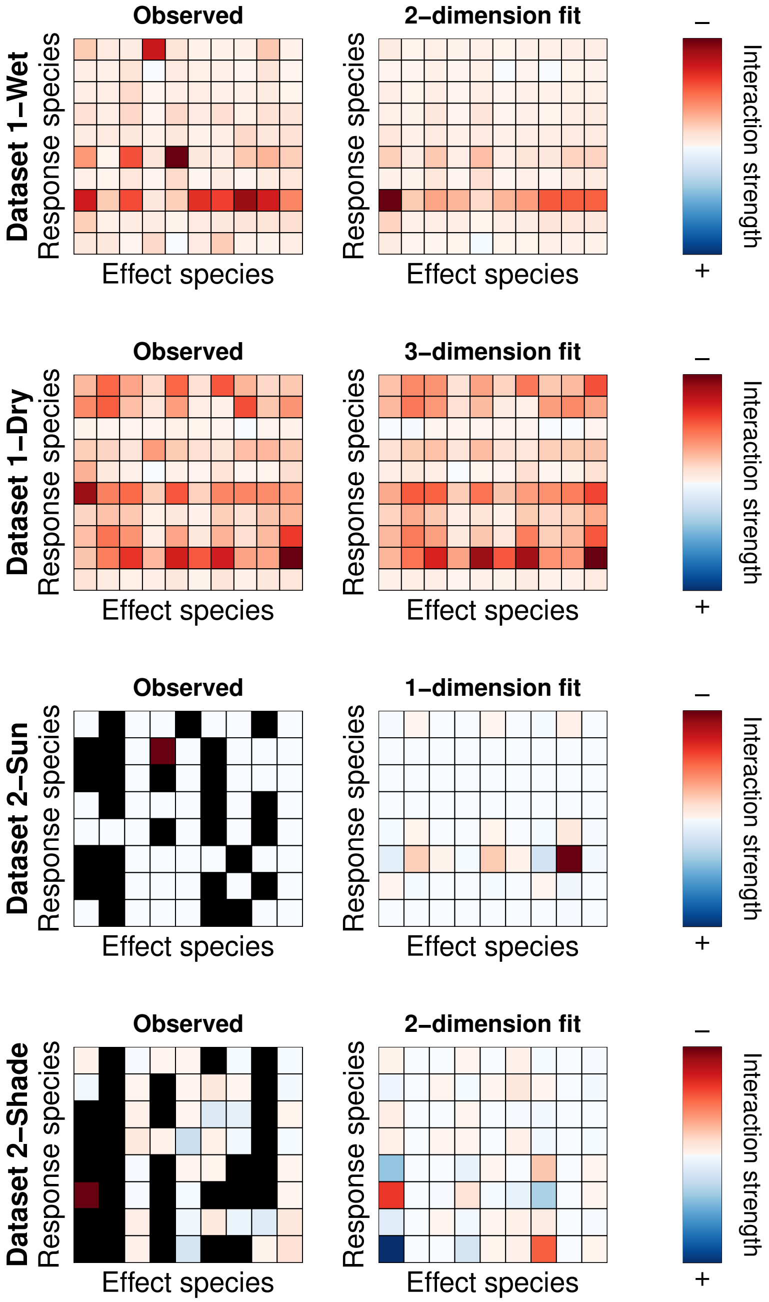


**Figure S3:** On the left of each row, we show the observed interaction matrix for each dataset. To the right of the observed, we show the inferred interaction matrix at niche dimensionality $\hat{d}$ as noted in Table S1. Interaction strengths have been scaled by dataset such that the most competitive is red, the most facilitative is blue, and no pairwise effect is white as noted in the colorbars on the far right.

### Observed versus fit interactions for all datasets (continued)

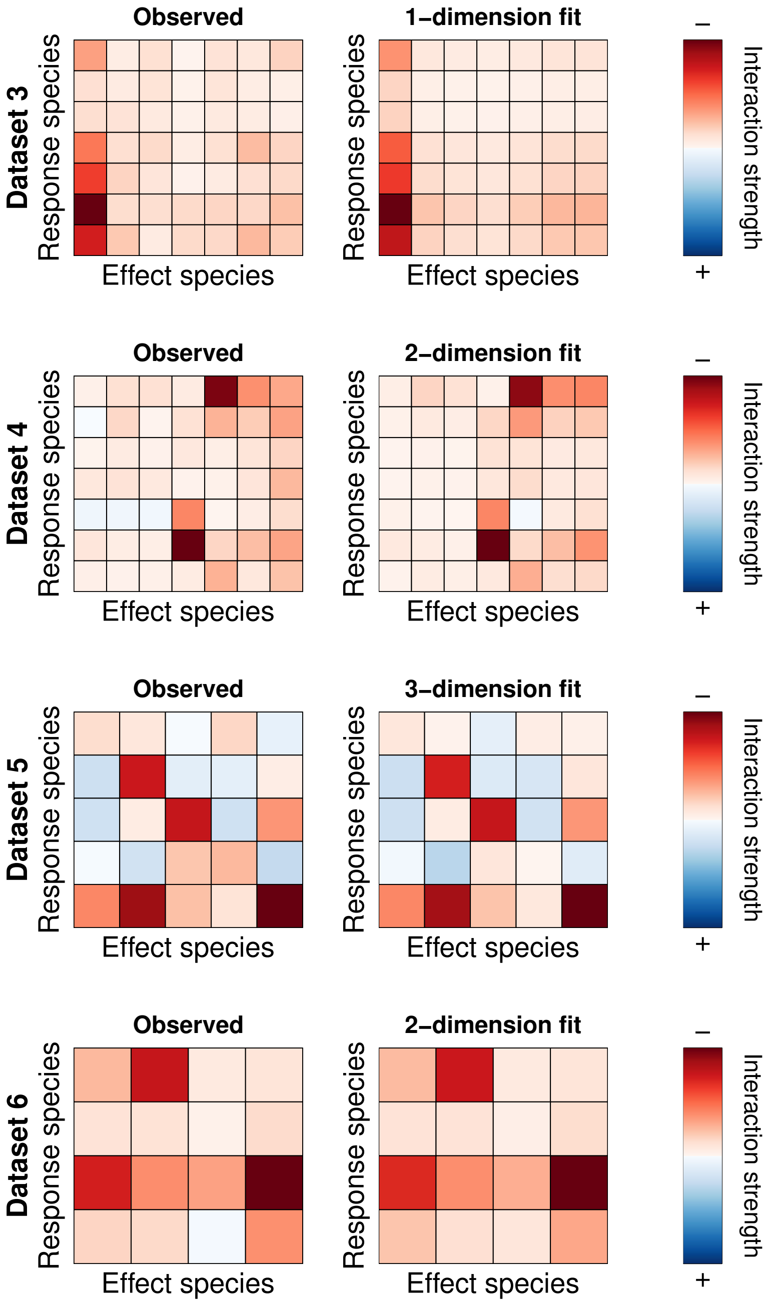

#### Observed versus fit interactions for all datasets (continued)

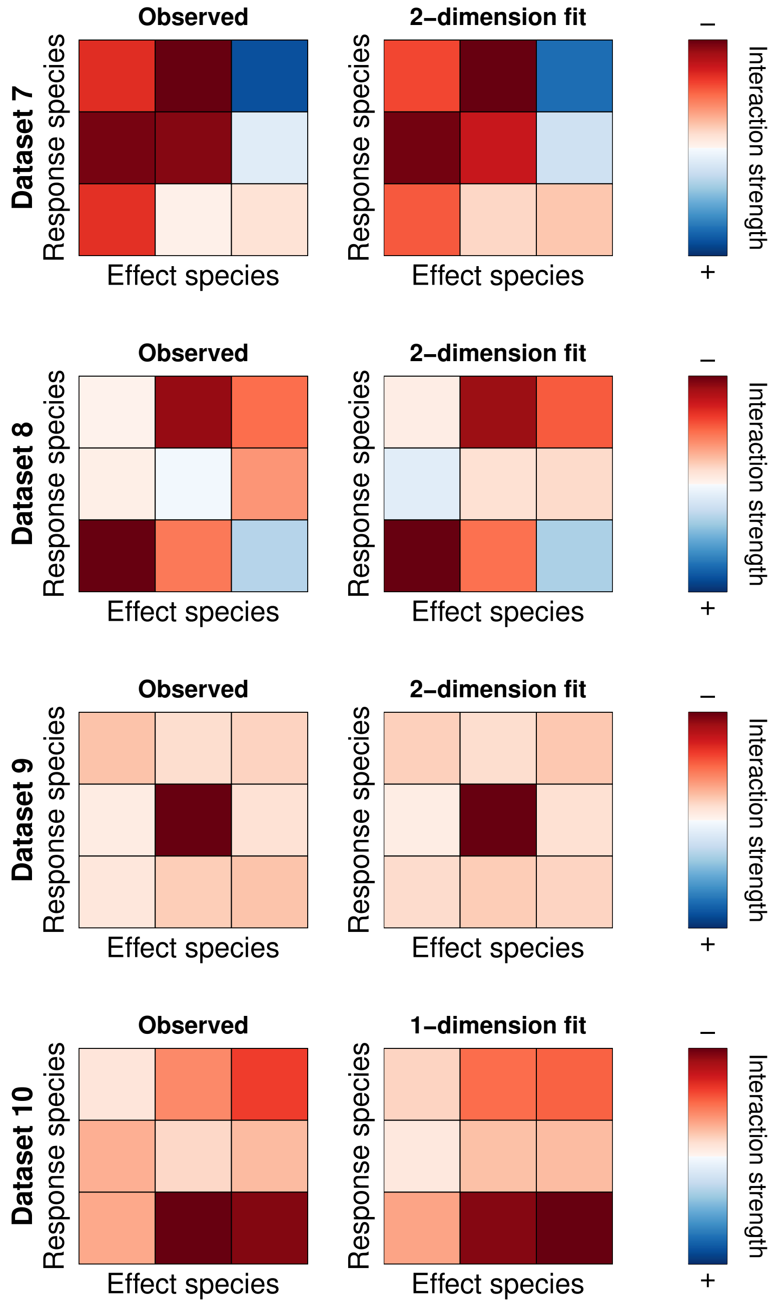

#### Observed versus fit interactions for all datasets (continued)

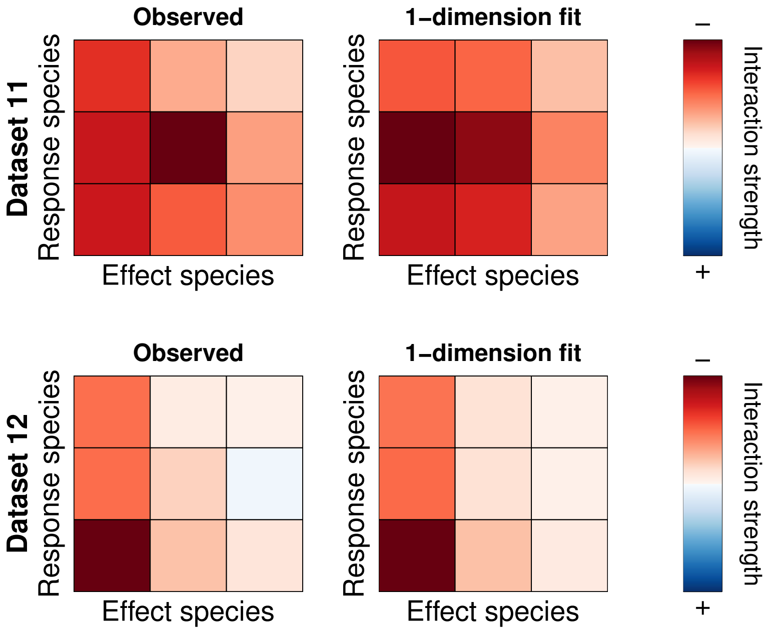

#### Response and effect traits for all datasets

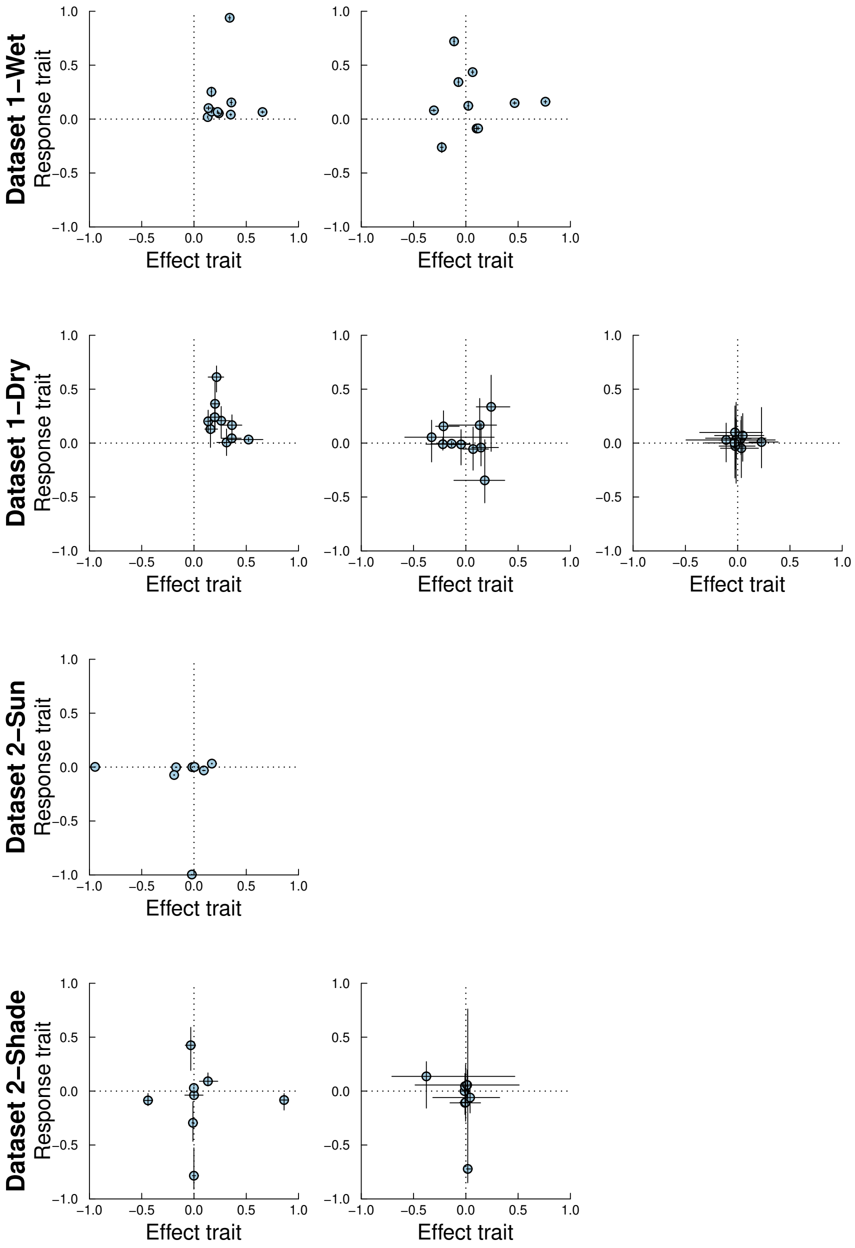


**Figure S4:** In each row, we show the relative position of all species in a dataset in response–effect trait space as in Fig. 3 for the first trait dimension of Dataset 1-Wet and Dataset 1-Dry. Recall that the number of response and effect traits for each species is equivalent to the inferred niche dimensionality $\hat{d}$ of the assemblage. Therefore, some datasets (rows) necessarily have fewer panels than others. Likewise, matrix-only datasets lead to deterministic response and effect traits and hence have no associated standard errors.

#### Response and effect traits for all datasets (continued)

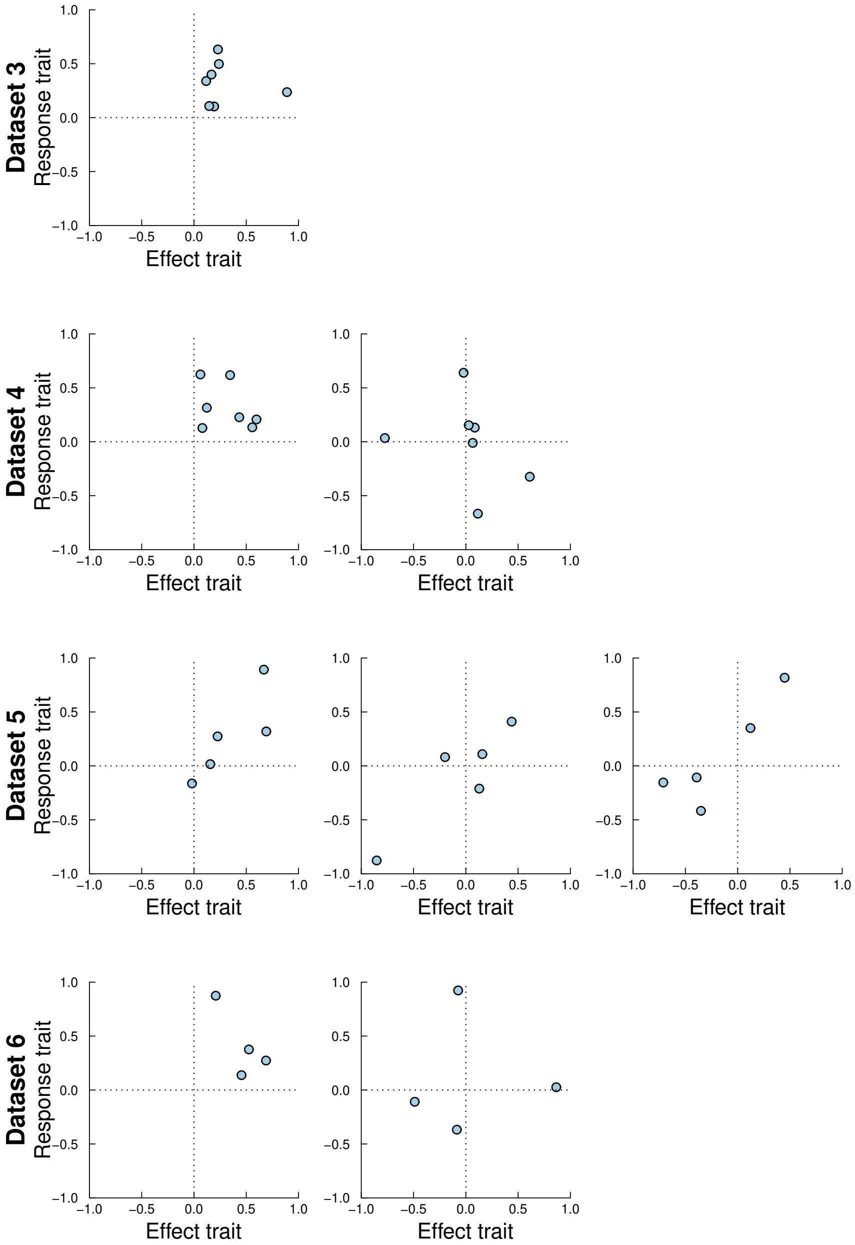

### Response and effect traits for all datasets (continued)

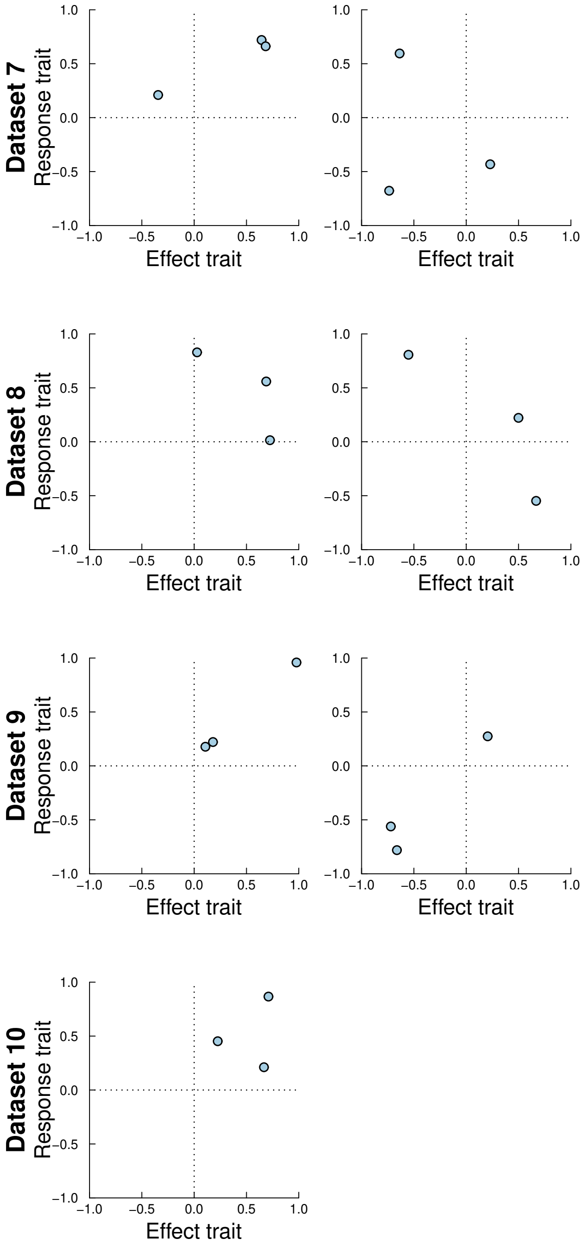

#### Response and effect traits for all datasets (continued)

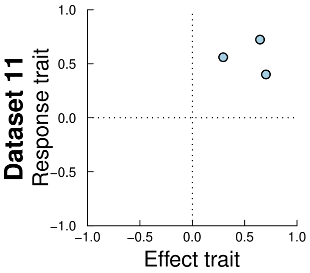


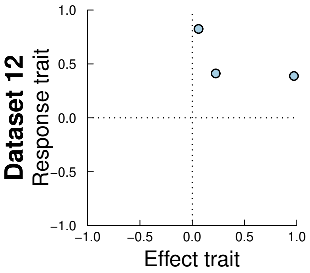

#### Niche dimensionality of randomised data

Just as random data may occasionally be perfectly correlated (or perfectly uncorrelated), we wished to ensure that our observation of low niche dimensionality across all empirical assemblages was not a spurious outcome of our methodology. To check this, we used Dataset 3 as a template from which to generate an ensemble of 25 randomized "equivalents" for which we could estimate $\hat{d}$. This dataset had 7 focal species, and we sampled their intrinsic growth rates $\lambda_i$ from a uniform distribution $U(0, 10)$ and their pairwise interaction coefficients $\alpha_{ij}$ from a uniform distribution $U(0, 1)$. We then simulated biomass production using Eq. (1) for each focal species growing with neighbour densities $N_j \in 1, 5, 10, 25, 50, 75, 100, 175, 250$—which is equivalent to the experimental conditions for the observed data[3]. We simulated one replicate observation for each focal–neighbour combination, giving a total of 441 simulated observations per randomized dataset. For each of these, we then estimated $\hat{d}$ based on the fit with lowest $AIC$. Our expectation was that $\hat{d}$ should tend to be equal to $S$ since the randomized data lacks a consistent underlying structure and/or competitive hierarchy. As expected, we found that the median $\hat{d} = 7$ (mode = 7; mean = 6.36; variance = 0.74) across these 25 randomized datasets (Fig. S5).

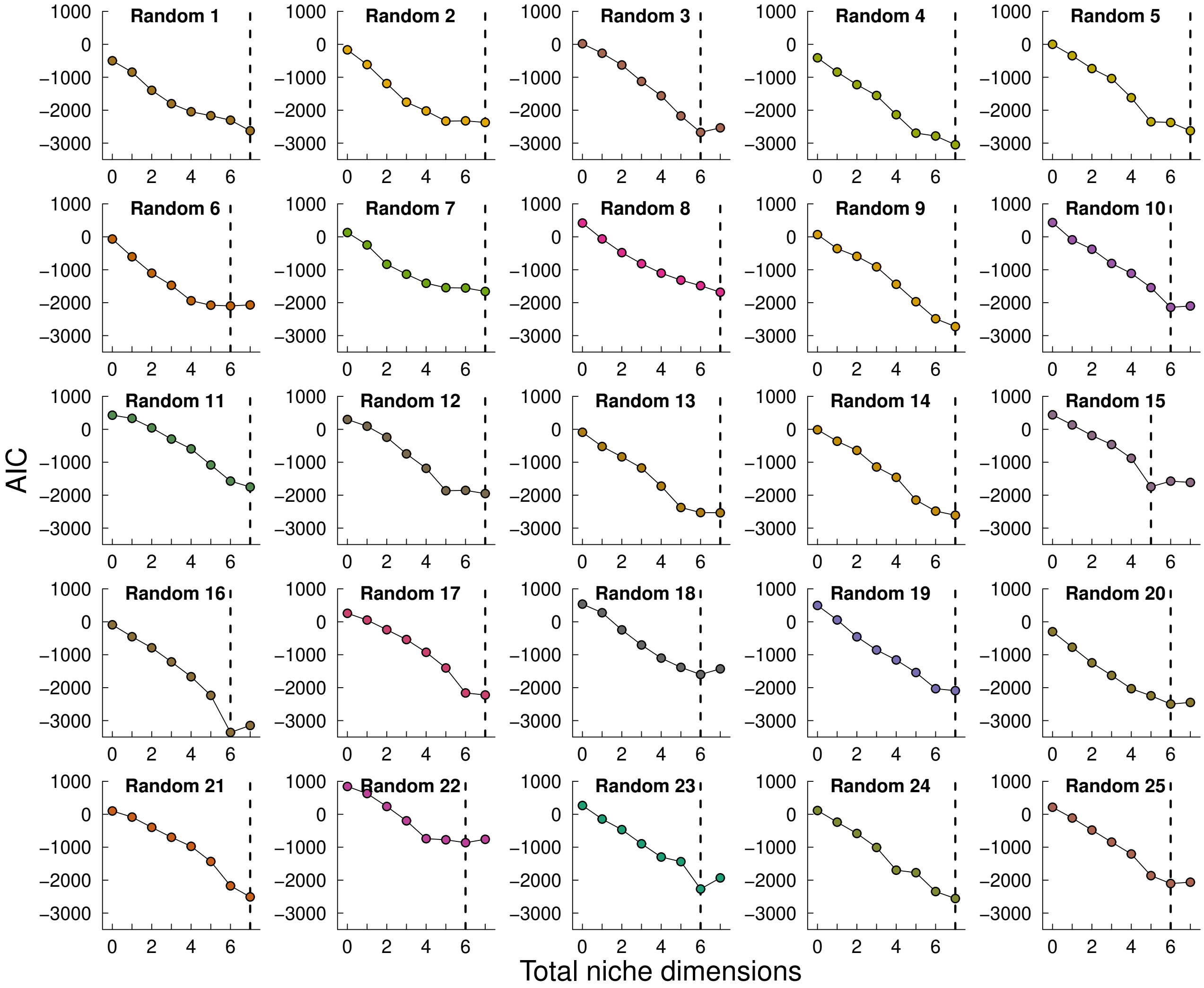


**Figure S5:** For an ensemble of randomly-generated data, we show the $AIC$ of model fits at fixed numbers of niche dimensions. The dashed line indicates the niche dimensionality $\hat{d}$ corresponding to minimum $AIC$.
